# Developing an *in silico* minimum inhibitory concentration panel test for *Klebsiella pneumoniae*

**DOI:** 10.1101/193797

**Authors:** Marcus Nguyen, Thomas Brettin, S. Wesley Long, Randall J. Olsen, James M. Musser, Robert Olson, Maulik Shukla, Rick L. Stevens, Fangfang Xia, Hyunseung Yoo, James J. Davis

## Abstract

Antimicrobial resistant infections are a serious public health threat worldwide. Whole genome sequencing approaches to rapidly identify pathogens and predict antibiotic resistance phenotypes are becoming more feasible and may offer a way to reduce clinical test turnaround times compared to conventional culture-based methods, and in turn, improve patient outcomes. In this study, we use whole genome sequence data from 1668 clinical isolates of *Klebsiella pneumoniae* to develop a XGBoost-based machine learning model that accurately predicts minimum inhibitory concentrations (MICs) for 20 antibiotics. The overall accuracy of the model, within ±1 two-fold dilution factor, is 92%. Individual accuracies are≥90% for 15/20 antibiotics. We show that the MICs predicted by the model correlate with known antimicrobial resistance genes. Importantly, the genome-wide approach described in this study offers a way to predict MICs for isolates without knowledge of the underlying gene content. This study shows that machine learning can be used to build a complete *in silico* MIC prediction panel for *K. pneumoniae* and provides a framework for building MIC prediction models for other pathogenic bacteria.

## Introduction

*Klebsiella pneumoniae* infections are a major cause of morbidity and mortality worldwide. Over the past several years, the emergence of antimicrobial resistant (AMR) *K. pneumoniae* strains has been increasing at an alarming rate, with reports of pan-resistant strains appearing in the literature and lay press^1–3^. Reports of hospital-based outbreaks are particularly concerning, and recent evidence suggests that AMR *K. pneumoniae* clones are circulating in the community^3–5^. As antimicrobial resistance increases, fewer effective antibiotics are available for physicians to treat these life-threatening infections. In response, the World Health Organization recently listed carbapenem and third generation cephalosporin resistant Enterobacteriaceae (including *K. pneumoniae*) among the most critical organisms for antimicrobial drug development^6^.

When a patient is diagnosed with an infection, it is critically important to prescribe appropriate antimicrobial therapy as quickly as possible. Rapid pathogen identification and appropriate antimicrobial therapy administration significantly decreases mortality, improves patient outcomes, reduces health care costs, and decreases the use of ineffective or inappropriate antibiotics^7–9^. For bloodstream infections, mortality increases every hour that appropriate therapy is delayed^9^. The conventional clinical microbiology laboratory evaluation of a suspected infection requires inoculation of the specimen on primary culture media and incubation until there is sufficient growth to perform taxonomic identification and minimal inhibitory concentration (MIC) determination. In many cases, subcultures are needed to purify mixed cultures containing more than one organism or generate sufficient colonial material for testing. Depending on the growth rate of the organism and the MIC testing procedures used, the multiple culture steps can add one or more days to the laboratory workup^10, 11^.

Compared to conventional culture-based methods, rapid molecular assays may significantly reduce turnaround times by eliminating one or more subculture steps. The most common sequence-based methods for predicting the AMR phenotypes of an organism identify the presence of genes implicated in resistance using PCR, microarrays or whole genome sequencing^12–14^. Well-designed gene-based detection methods are capable of providing an accurate prediction of susceptibility or resistance for the genes tested, but and there are several limitations to this approach. First, it relies on well-curated databases of AMR genes, which can be difficult to maintain^15–18.^ For example, the commonly used databases of AMR genes are currently excellent at cataloging well-studied AMR genes like *β*-lactamases^19^, but often lack data for diverse efflux mechanisms^18^. Second, similarity-based matching strategies to determine AMR gene content may incorrectly assign AMR functions to paralogous non-AMR genes. Gene based approaches may also fail to identify critical mutations in intergenic regions, including regulatory and promoter sequences, leading to a false-negative susceptibility prediction. Also, PCR based methods use primers for amplification, which may not anneal if mutations are present in the complementary region, again rendering an incorrect result. Finally, since these methods are based on preexisting knowledge of AMR conferring genomic regions, they are not able to predict resistance if the molecular mechanism is unknown or multifactorial.

The public sharing of whole genome sequence data with clinical AMR metadata has enabled the use of machine learning (ML) methods that predict AMR phenotypes without relying on a database of preexisting AMR genes or mutations. Two recent studies have used this approach to obtain accurate predictions of susceptibility or resistance in organisms with no *a priori* information about the gene content of the organisms^20,21^. To do this, they used short nucleotide k-mers as features and the laboratory derived AMR phenotypes as labels. Other studies have successfully used AMR genes, SNPs, and whole genome sequence data (or a combination thereof) to build ML classifiers with good accuracies^22–27^. Recent examples of gene-based and whole genome-based classification approaches for *Klebsiella* were reported by Stoesser et al.^27^, Long et al.^3^, and Pesesky et al.^24^.

To date, most AMR prediction methods have focused on classifying “susceptible” and “resistant” phenotypes. While simple and oftentimes sufficient, this approach can be error prone because it relies on the clinical interpretations of break point values. Also, intermediate phenotypes do not fit within this classification scheme. A small number of studies have attempted to predict MICs based on gene content^28–30^. One notable recent publication used an ML algorithm trained on the SNPs from several key AMR genes to successfully predict MICs for *Neisseria gonorrhoeae*^29^.

In this study, we present an ML approach for predicting MICs for *K*. pneumoniae. Our strategy requires no *a priori* knowledge of the underlying gene content. The current model offers MIC prediction for 20 antibiotics. To our knowledge, this is the largest MIC prediction study for a human pathogen to date. We discuss the strengths and limitations of our approach and the necessary steps required to implement *in silico* MIC prediction using whole genome sequence data for *K. pneumoniae* in the clinical laboratory.

## Results

### Approach

For several years, the microbiology laboratory at Houston Methodist Hospital System has been banking clinical isolates of *K. pneumoniae*. We recently sequenced the genome of AMR *K. pneumoniae* strains recovered from patients between 2011 and 2015^3^. Our goal is to use whole genome sequencing to detect the emergence of highly virulent clones, monitor the spread of AMR, and guide patient care decisions^31, 32^. We routinely perform whole genome sequencing in our clinical laboratory. Importantly, as the cost and speed of whole genome sequencing continues to decrease, it increasingly becomes a viable option for routine microbial diagnostics.

Using the whole genome sequence data for our *K. pneumoniae* clinical isolates, we sought to build an ML model that accurately predicts the MIC for 20 antibiotics. We chose a strategy that uses the entire genome as input, rather than individual genes, since this approach requires no *a priori* knowledge of the underlying gene content, and could potentially use data from uncharacterized AMR genes, intergenic or polymorphic coding regions, or non-AMR genes that may indirectly effect the MIC. To accomplish this, we computed the counts of all overlapping 10-mer oligonucleotide k-mers and combined them with the clinical laboratory generated MIC data for each antibiotic to form one large matrix containing both the k-mers and antibiotics as features. After exploring the problem as both a multiclass classification problem and a regression problem and evaluating common ML algorithms, we chose an extreme gradient boosting regression model through the XGBoost library^33^. We then iteratively evaluated the available parameters of the algorithm to maximize the accuracy of the model (Figure 1, Materials and Methods).

**Figure 1:**
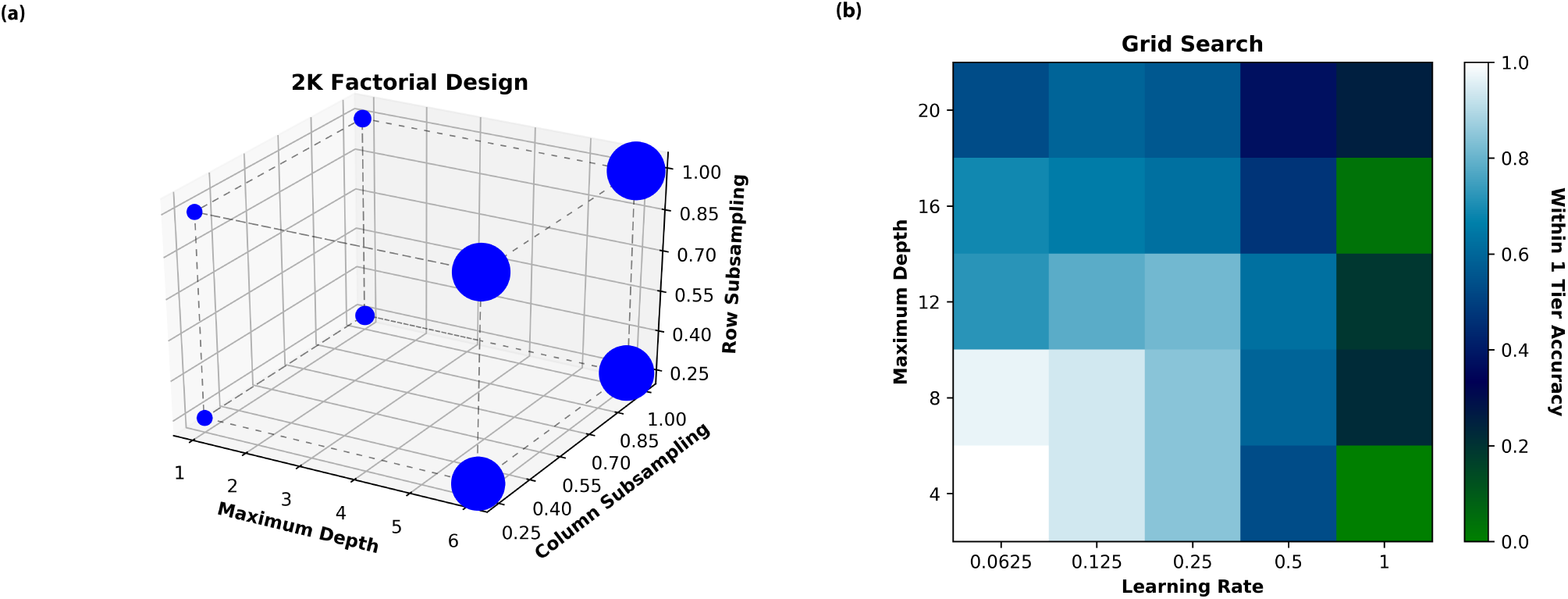
Results of the 2*^k^* factorial design on the XGBoost model. **(a)** Three dimensional plot showing the relationship between maximum depth, column subsampling and row subsampling parameters. The size of the spheres represent the within 1-tier accuracy score for the given model. Larger spheres indicate higher accuracy. Dashed lines are added to aid in visualization. The image shows that maximum tree depth plays the largest role in the XGBoost model with column and row subsampling having smaller roles. **(b)** Heat map showing the relationship between learning rate and maximum tree depth. In the color scheme, white is most accurate and dark blue and green are least accurate. The image shows that lower depth and learning rates produce more accurate models.

### Model Accuracy

A 10-fold cross validation was used to access the overall stability and accuracy of the model. The raw accuracy of the model, defined as the ability to predict the exact MIC for a given genome and antibiotic (Supplemental Table S1), and accuracy within ±1 two-fold dilution factor (or 1-tier) of the actual MIC was measured (Table 1). Bounding the accuracy to within one two-fold dilution factor of the laboratory determined MIC is consistent with current FDA standards for diagnostic tools and conventional clinical microbiology practices^34, 35^. We also evaluated the model based on the very major error (VME) rate, defined as a resistant isolate having a MIC that is predicted to be susceptible, and the major error rate (ME), defined as a susceptible isolate having a MIC that is predicted to be resistant. MIC thresholds for susceptibility and resistance for clinical data and model predictions are based on current CLSI breakpoints^36^

**Table 1:** Accuracies for the entire XGBoost model and for the individual antibiotics.

| Antibiotic | Samples | Accuracy<sup>a</sup> | 95% C.I.<sup>b</sup> |
| --- | --- | --- | --- |
| All | 32705 | 0.92 | 0.92, 0.92 |
| Amikacin | 1667 | 0.97 | 0.96, 0.98 |
| Ampicillin | 1666 | 1.00 | 0.99, 1.00 |
| Ampicillin/Sulbactam | 1664 | 0.99 | 0.99, 1.00 |
| Aztreonam | 1644 | 0.89 | 0.89, 0.90 |
| Cefazolin | 1667 | 0.96 | 0.95, 0.96 |
| Cefepime | 1571 | 0.61 | 0.58, 0.64 |
| Cefoxitin | 1645 | 0.90 | 0.89, 0.91 |
| Ceftazidime | 1667 | 0.92 | 0.91, 0.93 |
| Ceftriaxone | 1667 | 0.89 | 0.87, 0.90 |
| Cefuroxime sodium | 1575 | 0.99 | 0.99, 1.00 |
| Ciprofloxacin | 1664 | 0.98 | 0.97, 0.98 |
| Gentamicin | 1667 | 0.95 | 0.93, 0.96 |
| Imipenem | 1666 | 0.94 | 0.93, 0.95 |
| Levofloxacin | 1666 | 0.97 | 0.96, 0.97 |
| Meropenem | 1660 | 0.93 | 0.91, 0.95 |
| Nitrofurantoin | 895 | 0.96 | 0.95, 0.97 |
| Piperacillin/Tazobactam | 1662 | 0.78 | 0.77, 0.79 |
| Tetracycline | 1667 | 0.89 | 0.87, 0.90 |
| Tobramycin | 1666 | 0.95 | 0.94, 0.96 |
| Trimethoprim/Sulfamethoxazole | 1667 | 0.95 | 0.94, 0.96 |

<sup>a</sup> Average accuracy within $\pm 1$ two-fold dilution factor, based on a 10-fold cross validation.
<sup>b</sup> 95% confidence interval.

The average raw accuracy of the entire model, testing on all available MICs for 20 antibiotics was 69% with a 95% confidence interval of [68%; 69%]. The within 1-tier accuracy was much higher, at 92% with a 95% confidence interval of [92%; 92%] (Table 1, Supplemental Table S1). The large difference between the raw accuracy and the within 1-tier accuracy is probably the result of a variety of factors including the inherent error in the laboratory MIC testing procedure^37^, variations in growth and testing conditions, MICs with > and < values (which may actually represent a range of values), and a possible lack of discriminating genetic features between adjacent MIC dilutions. The raw accuracies and within 1-tier accuracies for the individual antibiotics track similarly, with the raw accuracies being lower and the within 1-tier accuracies being markedly higher. Overall, 15 of the 20 antibiotics have within 1-tier accuracies ≥90%. Three antibiotics have within 1-tier accuracies = 89%, while piperacillin/tazobactam and cefepime had lower within 1-tier accuracies of 78% and 61%, respectively (Table 1).

The accuracy of the model varies across the MICs for each antibiotic, in part, due to nonuniform representation of genomes for every possible value; however, the included genomes are representative of *K. pneumoniae* strains causing human infections. Figure 2 depicts the number of genomes and the within 1-tier accuracy for each MIC and antibiotic. Overall, MICs represented by many genomes tend to have high accuracies and MICs represented by fewer genomes tend to have lower accuracies. For example, the model has higher accuracies for *β*-lactam resistant MICs than susceptible MICs because there are fewer genomes for susceptible strains in our data set. In some cases, the model appears to be able to successfully interpolate over bins with a small number of samples. For example tobramycin MIC = 4*μ*g/ml, which is in between bins with deep sampling and high accuracy had 23 samples and an accuracy equal to 90%. Accuracies and confidence intervals for all bins are shown in Supplemental Table S2. The actual and predicted MICs for each genome are displayed in Supplemental Table S3.

**Figure 2:**
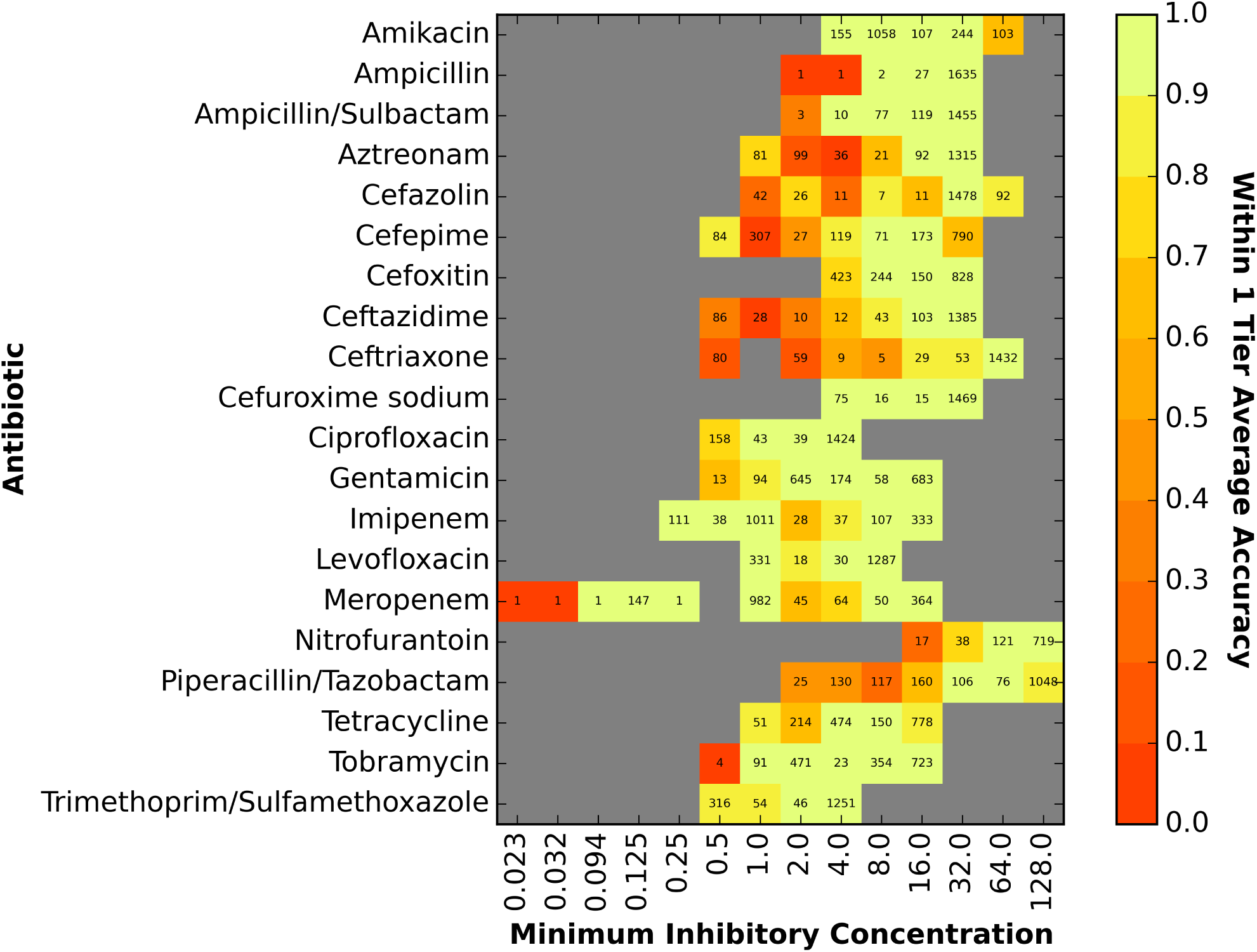
The accuracy of the XGBoost model for individual MICs. The X-axis of the heatmap shows the actual MIC (*μ*g/ml) for a bin and the Y-axis lists the antibiotics. The within ±1-tier accuracy of a particular antibiotic-MIC bin is denoted by color, with red and orange being least accurate and bright yellow and green being most accurate. The number within each cell represents the number of samples (genomes with the MIC) within the bin.

The average VME rate for the entire model was 3.1% with a 95% confidence interval of [2.8%;3.4%](Table 2). The average ME rate was 3.7% with a confidence interval of [3.3%;4.1%] (Table 2). The VME ranged from 0 for ampicillin and ceftriaxone to 30% for amikacin. Poorer prediction of amikacin MICs is likely due to the lower representation of amikacin resistant genomes in the dataset. Eleven of the antibiotics have VME rates with confidence intervals between 1.5 and 7.5% for the lower and upper bounds respectively. Likewise, 12 of the antibiotics have ME rates ≤ 3%. These results align with FDA diagnostic device standards^34^, suggesting that our model may be suitable for clinical use. However, it must be noted that susceptible and resistant MICs are not balanced in the data set, so the lowest VME rates tend to track with antibiotics that have the largest number of resistant genomes. We plan to test additional *K. pneumoniae* strains with these AMR phenotypes to fill this gap.

**Table 2:** Error rates for the entire XGBoost model for the individual antibiotics.

| Antibiotic | Resistant | Susceptible | VME<sup>a</sup> | VME 95% C. I.<sup>b</sup> | ME<sup>c</sup> | ME 95% C. I.<sup>b</sup> |
| --- | --- | --- | --- | --- | --- | --- |
| All | 21404 | 9410 | 0.031 | 0.028, 0.034 | 0.037 | 0.033, 0.041 |
| Amikacin | 103 | 1320 | 0.298 | 0.239, 0.358 | 0.000 | 0.000, 0.000 |
| Ampicillin | 1635 | 4 | 0.000 | 0.000, 0.000 | 0.000 | 0.000, 0.000 |
| Ampicillin/Sulbactam | 1455 | 90 | 0.003 | 0.000, 0.007 | 0.032 | −0.021, 0.085 |
| Aztreonam | 1407 | 216 | 0.001 | −0.001, 0.002 | 0.398 | 0.333, 0.462 |
| Cefazolin | 1570 | 97 | 0.060 | 0.047, 0.072 | 0.018 | −0.009, 0.046 |
| Cefepime | 963 | 418 | 0.007 | 0.002, 0.012 | 0.137 | 0.077, 0.197 |
| Cefoxitin | 828 | 667 | 0.077 | 0.060, 0.095 | 0.009 | −0.001, 0.019 |
| Ceftazidime | 1488 | 136 | 0.005 | 0.001, 0.008 | 0.123 | 0.069, 0.177 |
| Ceftriaxone | 1528 | 80 | 0.000 | 0.000, 0.000 | 0.188 | 0.101, 0.274 |
| Cefuroxime sodium | 1469 | 91 | 0.002 | 0.000, 0.004 | 0.010 | −0.013, 0.033 |
| Ciprofloxacin | 1424 | 201 | 0.005 | 0.000, 0.010 | 0.025 | 0.000, 0.050 |
| Gentamicin | 683 | 926 | 0.072 | 0.061, 0.082 | 0.009 | 0.001, 0.017 |
| Imipenem | 478 | 1160 | 0.040 | 0.012, 0.067 | 0.032 | 0.021, 0.043 |
| Levofloxacin | 1287 | 349 | 0.016 | 0.006, 0.025 | 0.020 | 0.006, 0.034 |
| Meropenem | 481 | 1134 | 0.048 | 0.034, 0.062 | 0.027 | 0.017, 0.038 |
| Nitrofurantoin | 719 | 55 | 0.018 | 0.009, 0.027 | 0.227 | 0.098, 0.356 |
| Piperacillin/Tazobactam | 1048 | 432 | 0.025 | 0.011, 0.038 | 0.012 | 0.000, 0.023 |
| Tetracycline | 778 | 739 | 0.114 | 0.095, 0.134 | 0.008 | 0.001, 0.015 |
| Tobramycin | 723 | 589 | 0.040 | 0.023, 0.057 | 0.012 | 0.002, 0.022 |
| Trimethoprim/Sulfamethoxazole | 1251 | 416 | 0.119 | 0.098, 0.140 | 0.108 | 0.082, 0.134 |

<sup>a</sup> VME, Average very major error rate defined as the percentage of resistant samples predicted as being susceptible.
<sup>b</sup> 95% confidence interval for the average VME and ME rates, respectively.
<sup>c</sup> ME, Average major error rate defined as the percentage of susceptible samples predicted as being resistant.

### Correlation with AMR Genes

The XGBoost algorithm generates a regression model that is based on an ensemble of decision trees. In our model, these decision trees split the data based on k-mers that distinguish the different antibiotics and MICs. Interpreting the features that are being selected by the model and subsequently understanding their relationship to a given antibiotic and resistance mechanism can be challenging for several reasons. Primarily, understanding the feature importance is an open graph theory problem. Furthermore, if an important k-mer maps to an uncharacterized gene or genomic region, it becomes difficult to determine if it is a hallmark of resistance or susceptibility. Finally, unambiguously associating a k-mer to a given antibiotic may inappropriately confer resistance to several related antibiotics, and we know that this dataset contains multiple resistance gene types such as the Bla-KPC and TEM *β*-lactamases^3^.

In general, the AMR that is detected in *K. pneumoniae* strains isolated from patients is usually conferred by the acquisition of antibiotic resistance genes rather than an accumulation of SNPs^3, 38^. We reasoned that in many cases, the MICs for an antibiotic should correlate with the occurrence of genes known to confer resistance to that antibiotic. By comparing the correlation of MICs and functional roles with the correlation that is based on the predicted MICs from the model, we gain an understanding of the relationship between MICs and AMR genes. We can also observe how closely the model is replicating these relationships. To do this, we computed the Pearson correlation coefficient (PCC) between the MICs for each genome and the presence or absence of each unique functional role in each genome (Table 3).

**Table 3:** The functional role that is most highly correlated with the MICs for each antibiotic.

| Antibiotic | PATRIC Functional Role | PCC Actual MIC<sup>a</sup> | PCC Predicted MIC<sup>b</sup> | Top 10 Coverage<sup>c</sup> |
| --- | --- | --- | --- | --- |
| Meropenem | Class A beta-lactamase (EC 3.5.2.6) => KPC family, carbapenem-hydrolyzing | 0.923 | 0.814 | 0.7 |
| Trimethoprim Sulfamethoxazole | Dihydropteroate synthase type-2 (EC 2.5.1.15) @ Sulfonamide resistance protein | 0.919 | 0.758 | 0.9 |
| Imipenem | Class A beta-lactamase (EC 3.5.2.6) => KPC family, carbapenem-hydrolyzing | 0.891 | 0.905 | 0.8 |
| Cefepime | Class A beta-lactamase (EC 3.5.2.6) => CTX-M family, extended-spectrum | 0.848 | 0.648 | 0.9 |
| Tobramycin | Aminoglycoside N(6')-acetyltransferase (EC 2.3.1.82) => AAC(6')-Ib/AAC(6')-II | 0.837 | 0.853 | 0.8 |
| Tetracycline | Tetracycline resistance regulatory protein TetR | 0.829 | 0.717 | 0.8 |
| Ceftriaxone | Class A beta-lactamase (EC 3.5.2.6) => CTX-M family, extended-spectrum | 0.823 | 0.700 | 0.7 |
| Gentamicin | Aminoglycoside N(3)-acetyltransferase (EC 2.3.1.81) => AAC(3)-II,III,IV,VI,VIII,IX,X | 0.818 | 0.862 | 0.6 |
| Ampicillin Sulbactam | Class A beta-lactamase (EC 3.5.2.6) => TEM family | 0.780 | 0.787 | 0.8 |
| Ciprofloxacin | Integron integrase IntI1 | 0.715 | 0.713 | 0.8 |
| Aztreonam | Integron integrase IntI1 | 0.678 | 0.614 | 0.7 |
| Cefazolin | Class A beta-lactamase (EC 3.5.2.6) => CTX-M family, extended-spectrum | 0.676 | 0.667 | 0.9 |
| Cefuroxime sodium | Aminoglycoside N(3)-acetyltransferase (EC 2.3.1.81) => AAC(3)-II,III,IV,VI,VIII,IX,X | 0.668 | 0.616 | 0.3 |
| Ceftazidime | Integron integrase IntI1 | 0.657 | 0.623 | 0.6 |
| Levofloxacin | probable bacteriophage protein STY1063 | 0.588 | 0.584 | 0.7 |
| Piperacillin Tazobactam | plasmid stabilization system | 0.583 | 0.501 | 0.1 |
| Amikacin | IncI1 plasmid conjugative transfer prepilin PilS | 0.577 | 0.478 | 0.2 |
| Cefoxitin | Class A beta-lactamase (EC 3.5.2.6) => KPC family, carbapenem-hydrolyzing | 0.550 | 0.571 | 0.6 |
| Nitrofurantoin | Integron integrase IntI1 | 0.433 | 0.507 | 0.6 |
| Ampicillin | Class A beta-lactamase (EC 3.5.2.6) => TEM family | 0.357 | 0.327 | 0.0 |

<sup>a</sup> Pearson correlation coefficient between the occurrences of the given functional role and the actual MICs.
<sup>b</sup> Pearson correlation coefficient between the occurrences of the given functional role and the predicted MICs.
<sup>c</sup> The fraction of the top 10 functional roles (by PCC) for the predicted MICs that occur in the top 10 for the actual MICs.

In the case of 12 antibiotics, we observe high PCCs between the MICs and the functional roles for well-known AMR genes.

For example, the *β*-lactam antibiotics correlate with the CTX, KPC and TEM type *β*-lactamase genes and the aminoglycoside antibiotics correlate with aminoglycoside acetyltransferase genes. For 8 of the 20 antibiotics, the association between the functional role and AMR is not obvious. In some of these cases, the functional roles appear to be related to horizontal gene transfer, and this may have resulted in their high PCCs. In other cases, the genes may not be sufficiently characterized by the raw 10-mer counts. When the analysis is repeated using MICs that are predicted by the model, the PCCs track very closely with those generated from the actual MICs, indicating that the model is not only learning that these genes exist (the model is only fed 10-mer counts, not whole genes), but it is also placing importance on these genes. Similarly, there is considerable overlap between the top ten most highly correlated functional roles based on the actual and predicted MICs (Table 3, last column).

In previous work, we built 16 AdaBoost-based classifiers for predicting susceptibility and resistance and installed them in the RAST annotation system^3^. Thirteen of the antibiotics that were used to build these classifiers are represented in our MIC prediction model. The top genomic regions predicted by the *K. pneumoniae* AdaBoost classifiers correspond to the most most highly correlated functional roles from the MIC prediction model in for four of the thirteen antibiotics. These include gentamicin, imipenem, meropenem and tetracycline. The top AdaBoost feature can be found found among the top five most highly correlated functional roles in three more cases including aztreonam, cefoxitin, and trimethoprim/sulfamethoxazole. Since the AdaBoost classifiers match SNPs in gyrase and topoisomerase genes conferring resistance to ciprofloxacin and levofloxacin, assessing the correlation to the presence of functional roles will not work for these two antibiotics. It is not immediately clear why the top features for the other four antibiotics (amikacin, cefepime, piperacillin/tazobactam and tobramicin) are not highly correlated, but no characterized AMR roles appear in the top 10 most highly correlated features.

### Building a Model Based on AMR Genes

Although we chose to build a model that was based on data from the entire genome, previous studies have built MIC prediction models using the known AMR genes ^28–30^. We wanted to know whether a model based on whole genome data, or a model based on only AMR genes would be more accurate. On one hand, the extra k-mers used by the whole genome model could be a source of noise, but on the other hand, they may represent useful data that could making the model more accurate. To make this comparison, we built a model that used only the AMR genes as the source of k-mers, keeping all other parameters identical. Both the PATRIC annotation service ^18, 39^ and the CARD database^15^ were used as sources of potential AMR genes. The overall accuracies for the whole genome and AMR gene-based models are both 92%. Likewise, the individual accuracies for each antibiotic are also nearly identical, differing by no more than 2% for any antibiotic (Table S4, Figure S1). This indicates that the extra k-mers used by the whole genome are not a source of noise for the XGBoost model. Since the AMR genes model is much smaller than the whole-genome model (20GB vs. 148GB), it is more efficient to compute, and it is therefore tempting to conclude that a model built from AMR genes is sufficient for performing MIC prediction for *K. pneumoniae*. However, due to the smaller number of susceptible and intermediate samples in this data set, more genome sequencing is necessary to determine if a model built from AMR genes is sufficient for accurately predicting these lower MIC values.

When we repeat this analysis building an identical model for the leftover non-AMR genes, we also observe an overall accuracy of 92%, and accuracies for individual antibiotics that track closely with the models built from full contigs and AMR genes (Table S4, Figure S1). This indicates that there is enough residual data in the non-AMR genes to accurately predict MIC values as well. This could be due to the presence of uncharacterized AMR genes in the data set, the existence of relevant information such as non-AMR genes co-occurring with AMR genes, or variants in non-AMR genes that have an impact on MICs. In the case of all three models, it is unlikely that the accuracy is due to the model mapping to a phylogenetic pattern between strains, since nearly clonal strains of the same MLST type can have a variety of different MIC values per antibiotic (Table S5).

## Discussion

Using the clinical laboratory determined MICs and whole genome sequence data for 1668 *K. pneumoniae* strain recovered from infected patients, we built an XGBoost regression model that can predict the MICs for 20 antibiotics with an average within ± 1-two fold dilution factor accuracy of 92%. Individually, 15 of the 20 antibiotics have within 1-tier accuracies >90%. These results demonstrate that accurate genome-based MIC prediction is possible for *K. pneumoniae* isolates. Herein, we provide the necessary groundwork for building a complete *in silico* panel.

To date, our whole genome sequencing efforts have focused on the most highly antibiotic resistant pathogens, including extended spectrum *β*-lactamase-(ESBL) producing *K. pneumoniae*^3^, so the model currently lacks sufficient inclusion of organisms with MICs in the lower range. We also currently lack sufficient data to predict MICs for some less-commonly tested antibiotics, including amoxicillin/clavulanate, ertapenem, fosfomycin, moxifloxacin ticarcillin/clavulanate and tigecycline. Ideally, selecting a balanced number of organisms at each possible MIC could improve the overall accuracy of the model. Our future efforts will seek to close these gaps. Inclusion of additional strains will also improve the ME and VME rates. The *K. pneumoniae* strains used in our model were collected as part of a large, comprehensive, population-based study in Houston. Although our sampling capacity is extensive and we treat a diverse population of patients from Houston and around the world, the model may be further improved by including samples from other geographic locations.

In previous work, we built binary classifiers that can predict susceptibility or resistance for a given species and antibiotic^18, 20^.Although somewhat simplistic in approach, the method provides a straightforward way to extract the genomic features relating to resistance. In this study, in order to achieve high accuracies for predicting each MIC, we combined the antibiotics and k-mers into a large matrix and used XGBoost as the ML method. While the approach is clearly advantageous because it provides a more refined phenotype prediction, feature extraction from this kind of model remains quite challenging. Although we have shown strong correlations between the actual MICs and predicted MICs with known AMR genes, we will continue to explore ways to extract the key gene data that contributes to each MIC. Importantly, these studies may provide new insight to the molecular mechanisms conferring intermediate phenotypes.

Using data from the entire genome is advantageous for building the MIC prediction model because it requires no underlying information about the gene content of the organisms. However, the current data set lacks sufficient sampling of genomes with susceptible and intermediate phenotypes to determine if models built from whole genome data or only the AMR genes will ultimately differ in accuracy. Since genomic features that are not annotated as being directly involved in conferring AMR are likely to influence on lower MIC values, a whole-genome approach might be beneficial for discriminating low MIC values as we acquire more data. Surprisingly, we found that there is sufficient data to perform accurate MIC prediction using only the non-AMR genes. This may have value for building models that can predict AMR phenotypes from incomplete sequence data, as may be the case with a metagenomic samples. It may also provide a means by which to explore the more subtle effects that non-AMR genes may be having on AMR phenotypes.

In this study, we found that deeper trees, with a depths of 3–4, were optimal for the XGBoost model. A logical next step will be to train a deeper model, like a neural network, on this data set to determine if the accuracy can be further improved. A deep learning approach may also provide more efficient memory usage and reduced computational times. However, this strategy would not improve the extraction and interpretation of AMR-related genomic features, since feature extraction from deep learning methods is more challenging compared to ensemble methods such as XGBoost.

The genomes used in this study were sequenced using Illumina sequencing technology. In order to generate genome sequence data cost effectively, we collected samples and performed highly multiplexed runs in batches. However, this is not a clinically time efficient strategy. Newer devices such as the PacBio Sequel and Oxford MinION could potentially be used for point of care sequencing, and thus, may become a model for whole genome sequence-based diagnostic strategies^40^. However, at present, the cost of sequencing individual pathogens using these technologies is higher, and their error rates may be too high for effective MIC prediction with our ML method^41^. In order to couple the MIC prediction model outlined in this study with these sequencing platforms, we may need to either incorporate an error correction model for processing MinION or PacBio reads, or regenerate the model using genomes sequenced with these technologies. Further analyses to sequence based off of blood enrichment cultures, or from the actual wound source, rather than the pure culture, would also provide more rapid results, but require algorithms for identifying pathogens and eliminating host DNA and other contaminants. Regardless, sequence-based MIC prediction appears to be feasible as a diagnostic strategy.

## Methods

### Strain collection

*Klebsiella pneumoniae* isolates were cultured from patient specimens in the Houston Methodist Hospital System between September of 2011 and March of 2017. Strains were tested for minimum inhibitory concentrations to 20 antibiotics using the BD-Phoenix system (BD Diagnostics, Sparks, MD, USA). All of the strains collected before 2015 were originally part of two studies by Long and colleagues designed to track extended spectrum *β*-lactamase-(ESBL) producing strains ^3, 42^. A total of 1497 strains from the Long et al. study with BD-Phoenix MIC data were used in this analysis (Table S3). An additional 171 isolates, 93 ESBL-producing and 78 non-ESBL-producing (Phoenix ESBL test; BD), were also panel tested, whole genome sequenced, and used in this study. In total, we analyzed 1668 *Klebsiella pneumoniae* genomes.

### DNA extraction and whole-genome sequencing

Genomic DNA was extracted using the manufacturer’s Gram-negative protocol for the DNeasy Blood and Tissue kit (Qiagen) and quantified using a Qubit 3.0 instrument (Invitrogen). Whole-genome sequencing libraries were prepared using NexteraXT reagents (Illumina) and sequenced on a MiSeq or NextSeq 500 instrument (Illumina).

### Data preparation

When the MICs produced from the BD-Phoenix tests exceeded the testing thresholds, they were cleaned to remove the *>, <, ≥*, and *≤*symbols. If the MIC was *> x*, the MIC label was changed to 2*x*, if it was *< x*, the MIC was changed to 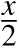, and if the MIC was *≥ x* or *≤ x*, the symbol was removed and the MIC remained unchanged. The *Log*_2_ value of these cleaned MICs were used for all ML tasks. For dual antibiotics with two MICs, such as trimethoprim/sulfamethoxazole, the first value was used in all cases, since the second value is either constant or dependent on the first.

### Experimental design

Genomes were assembled with SPAdes^43^ using the PATRIC assembly service^39^. Contigs with ≤ 500*bp* and ≤ 5-fold coverage were removed. The contigs were divided into overlapping 10-mer oligonucleotide k-mers, sorted in lexicographical order and counted using the software package KMC2^44^. K-mer counts and antibiotic names were used as input features into the model. Antibiotic names were one-hot encoded. In this study, we chose to use 10-mers as features, rather than a longer k-mer length, in order to reduce the size of the matrix. This enabled us to load the matrix into memory and perform cross validation on a machine with 1.5TB RAM. The dataset was then split into equal training, validation and testing sets. Each subset was created so that it contained the same number of MICs for a given antibiotic. The final distribution antibiotics used in the model is uniform, but the number of samples for a given MIC per antibiotic was not uniformly distributed.

The prediction of MICs can be cast as a regression problem or a multi-class classification problem with each MIC representing an individual class. We explored several popular ML algorithms including AdaBoost (Adaptive Boosting)^45^, bagging^46^, random forest^47^, extremely randomized trees^48^, and support vector machines^49^ from the scikit-learn python package^50^, as well as XGBoost (Extreme Gradient Boosting)^33^. Using default parameter choices, we attempted to cast the problem as both a classification and regression problem depending on the capabilities of the algorithm. Algorithms were compared based on accuracy and computational resources required. An XGBoost regression model was ultimately chosen for this study because it produced the highest default accuracies and had reasonable computation costs for the current data set.

To assess the sensitivity of the model with regard to the training data, we performed a ten-fold cross validation by taking all samples and randomly splitting them into 10 mutually exclusive sets. Each set was split such that all sets had an equal (or nearly equal) number of antibiotic-MIC combinations. Ten models were then generated using 8 of the sets for training, one for validation, and one for testing. In total, 10 models were created. The accuracy within ±1 two-fold dilution factor was computed along with the 95% confidence intervals for each model. This aligns with clinical practice and FDA device guidelines^34^.

### Hyperparameter tuning

Important hyperparameters from XGBoost were then selected and tuned using a 2*^k^* factorial design^51^ and a grid search, respectively. The model was tested for stability using a 10-fold cross validation. The 2*^k^* factorial design was used to tune 3 XGBoost parameters: maximum tree depth, column subsampling and row subsampling. The maximum tree depth parameter limits the maximum height of a decision tree while creating the ensemble. A value that is too high tends to overfit data whereas a value that is too low tends to underfit data^33^. A low value of 1 and a high value of 6 (6 is the default) were used to evaluate the impact of tree depth on the accuracy of the model. The column subsampling parameter limits the number of features that are chosen for training each tree in the ensemble. For example, if 0.5 is chosen, 50% of the features (k-mers) will be randomly chosen to train the first tree, then a different 50% for the second, and so on. A low value of 0.25 and a high value of 1 (1 is the default and maximum value allowed for XGBoost) were chosen. The row subsampling parameter limits the number of samples that are chosen for training for each tree in the ensemble. For example, if 0.5 is chosen, 50% of the samples (MIC tests) will be randomly chosen to train the first tree, then a different 50% for the second, etc. A low value of 0.25 and a high value of 1 (1 is the default and maximum value allowed for XGBoost) were chosen for evaluation.

The accuracy of the XGBoost model was evaluated in two ways. First, the accuracy of each individual MIC prediction over the test set was assessed within ± 1 two-fold dilution factor. Secondly the coefficient of determination, or R^2^, was also used as a metric during the hyperparameter tuning.

A 2*^k^* factorial design assumes that as a parameter is increased or decreased, the metric that is being tested will either increase or decrease. Additionally, it only gives an idea of the parameters that are important and not the optimal value for a given parameter. Since the accuracy is not known to always increase or decrease as the maximum tree depth value goes up^33^, this parameter needed to be tested systematically. A grid search^52^ was designed based on the results of the 2*^k^* factorial design to deal with both issues. Since learning rate is known to have a relationship with maximum tree depth, we systematically varied the maximum tree depth and the learning rate, in tandem, in a grid search experiment with respect to accuracy^33^.

The results of the 2*^k^* factorial design showed that when tuning the model, a higher maximum tree depth was preferable. Different values of row and column subsampling did not cause variance in the accuracy of the model, though a larger value was deemed to produce a more accurate model. Both factors had already been maximized at 1 during the 2*^k^* factorial. Figure 1a shows this in better detail.

The applied grid search showed that a lower value for the learning rate always returned a better solution. However, as the learning rate decreased, the training time increased. The gain in accuracy was deemed too small for the price (time) with a learning rate of 2*^-4^*. During testing we also found that a maximum depth between 3 and 4 was optimal for the *Klebsiella* data used to train the algorithm. Figure 1b shows this in better detail.

### Genome Annotation and AMR gene analyses

All genomes were annotated using the PATRIC annotation service in August of 2017^39^. MLST (Multi Locus Sequencing Typing) was performed by the PATRIC annotation service and was based on Diancourt et al. ^53^. To compare the model described above with one that was based only on AMR genes, we gathered the set of genes with AMR functional roles (annotations) defined by PATRIC^18^ as well as all genes encoding proteins matching the CARD database^15^ with BLASTP identity scores>= 80%^54^. The AMR gene-based model was computed the same as the final whole genome-based model described above. Non-AMR genes are defined as those that do not match the AMR set.

Correlation analyses were performed by first gathering the set of uniquely occurring functional roles from every *K. pneumoniae* genome. For each genome, the presence (+1) or absence (-1) of a functional role was compared to the MIC for an antibiotic. The Pearson correlation coefficient was computed for every functional role and antibiotic combination.

### Data Availability

The model and software for predicting MICs in for *K. pneumoniae* genomes can be found at the PATRIC github page: https://github.com/PATRIC3/mic_prediction. All genomes were submitted to the SRA under bioprojects (PRJNA376414, PRJNA386693 and PRJNA396774). SRA run accession numbers for individual genomes can be found in Table S3.

## Acknowledgements

This work was supported by the National Institute of Allergy and Infectious Diseases, National Institutes of Health, Department of Health and Human Service [Contract No. HHSN272201400027C] and the Fondren Foundation. We thank Concepcion Cantu, Matthew Ojeda Saavedra and Sarah Linson for technical assistance. We thank Emily Dietrich for her help preparing the manuscript.

## Author contributions statement

M.N., designed study, conducted experiments, performed data analysis, wrote paper; T.B., designed study; S.W.L., designed study, obtained strains, generated MIC data, performed data analysis, revised manuscript; J.M.M., designed study; R.O. conducted software engineering and distribution; R.J.O, designed study, obtained strains, generated MIC data, performed data analysis, revised manuscript; M.S., integration of data into PATRIC; R.L.S, designed study; F.X., designed study; H.Y. integration of data into PATRIC; J.J.D, designed study, performed data analysis, wrote paper. All authors reviewed the manuscript.

## Additional information

### Accession codes

All genomes were submitted to the SRA under bioprojects (PRJNA376414, PRJNA386693 and PR-JNA396774).

## Developing an *in silico* minimum inhibitory concentration panel test for *Klebsiella pneumoniae*

### Supplemental Information

**Table S1.** A comparison of raw accuracies and accuracies within ±1 two-fold dilution step of the actual MIC for the XGBoost model

| Antibiotic | Samples | Raw Accuracy<sup>a</sup> | Raw Accuracy<sup>a</sup><br>95% C.I.<sup>b</sup> | Within 1<br>two-fold<br>Accuracy<sup>c</sup> | Within 1<br>two-fold<sup>b</sup><br>95% C.I.<sup>b</sup> |
| --- | --- | --- | --- | --- | --- |
| All | 32705 | 0.69 | 0.68, 0.69 | 0.92 | 0.92, 0.92 |
| Amikacin | 1667 | 0.69 | 0.67, 0.71 | 0.97 | 0.96, 0.98 |
| Ampicillin | 1666 | 0.95 | 0.94, 0.96 | 1.00 | 0.99, 1.00 |
| Ampicillin/Sulbactam | 1664 | 0.88 | 0.86, 0.90 | 0.99 | 0.99, 1.00 |
| Aztreonam | 1644 | 0.62 | 0.6, 0.65 | 0.89 | 0.89, 0.90 |
| Cefazolin | 1667 | 0.85 | 0.84, 0.86 | 0.96 | 0.95, 0.96 |
| Cefepime | 1571 | 0.19 | 0.16, 0.22 | 0.61 | 0.58, 0.64 |
| Cefoxitin | 1645 | 0.40 | 0.37, 0.43 | 0.90 | 0.89, 0.91 |
| Ceftazidime | 1667 | 0.73 | 0.71, 0.76 | 0.92 | 0.91, 0.93 |
| Ceftriaxone | 1667 | 0.73 | 0.71, 0.75 | 0.89 | 0.87, 0.90 |
| Cefuroxime sodium | 1575 | 0.94 | 0.93, 0.95 | 0.99 | 0.99, 1.00 |
| Ciprofloxacin | 1664 | 0.87 | 0.85, 0.89 | 0.98 | 0.97, 0.98 |
| Gentamicin | 1667 | 0.65 | 0.62, 0.68 | 0.95 | 0.93, 0.96 |
| Imipenem | 1666 | 0.70 | 0.68, 0.71 | 0.94 | 0.93, 0.95 |
| Levofloxacin | 1666 | 0.82 | 0.80, 0.83 | 0.97 | 0.96, 0.97 |
| Meropenem | 1660 | 0.69 | 0.66, 0.71 | 0.93 | 0.91, 0.95 |
| Nitrofurantoin | 895 | 0.78 | 0.75, 0.82 | 0.96 | 0.95, 0.97 |
| Piperacillin/Tazobactam | 1662 | 0.46 | 0.43, 0.49 | 0.78 | 0.77, 0.79 |
| Tetracycline | 1667 | 0.50 | 0.48, 0.53 | 0.89 | 0.87, 0.90 |
| Tobramycin | 1666 | 0.61 | 0.57, 0.64 | 0.95 | 0.94, 0.96 |
| Trimethoprim/Sulfamethoxazole | 1667 | 0.74 | 0.72, 0.76 | 0.95 | 0.94, 0.96 |

<sup>a</sup> Average raw accuracy; raw accuracy is defined as predicting the actual MIC.
<sup>b</sup> 95% confidence interval.
<sup>c</sup> Average within $\pm 1$ two-fold dilution accuracy.

**Table S2.**
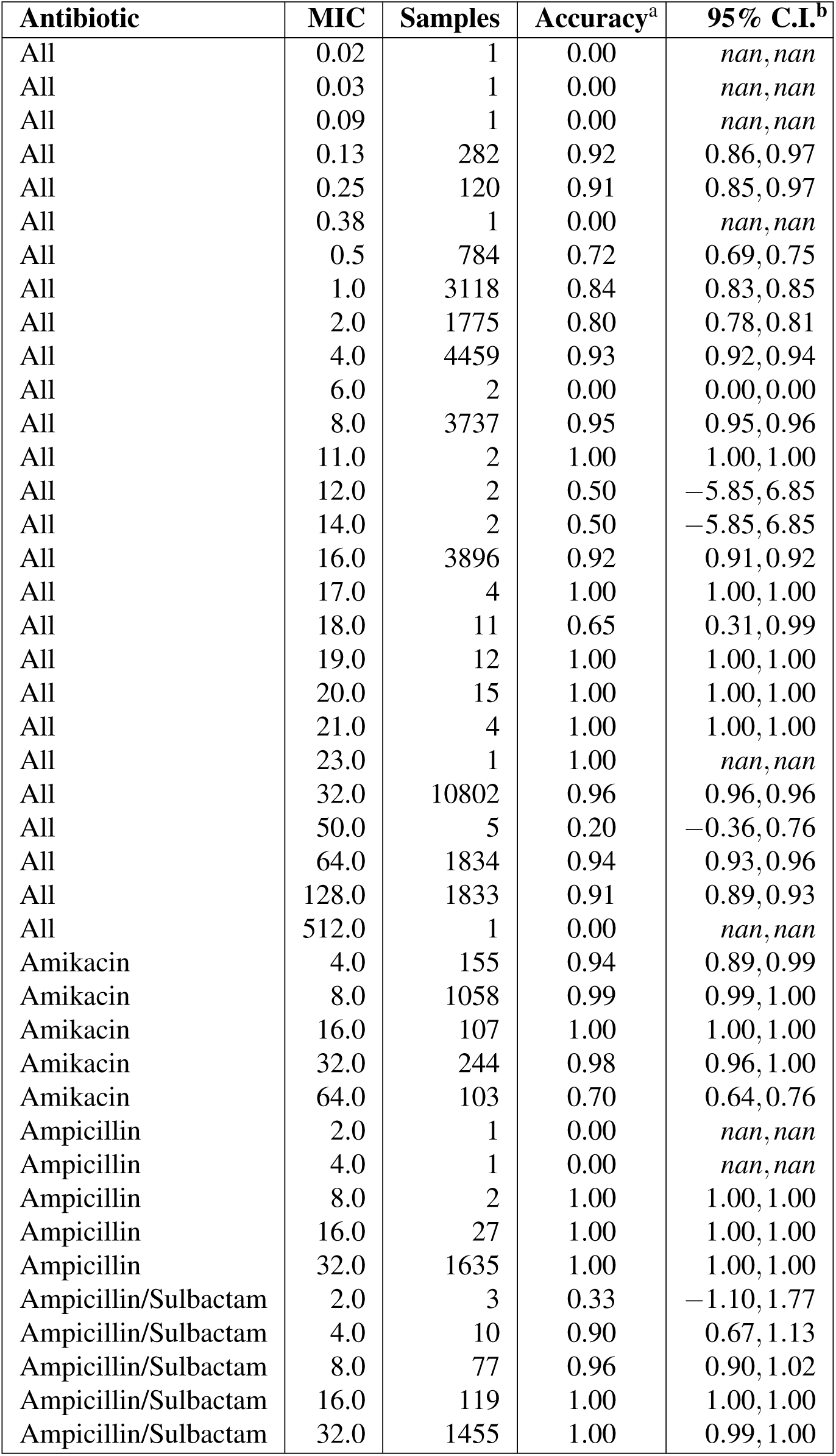
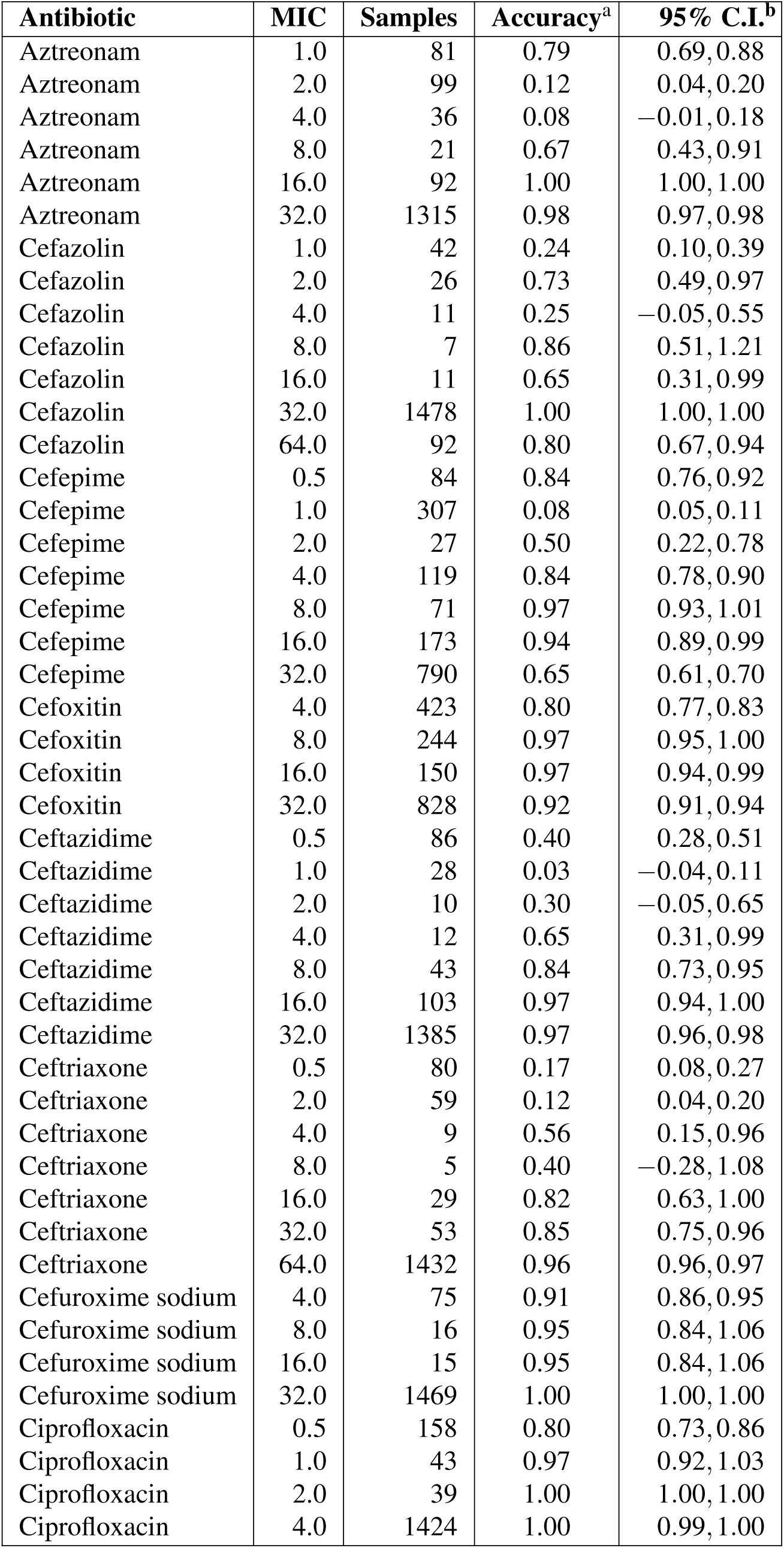
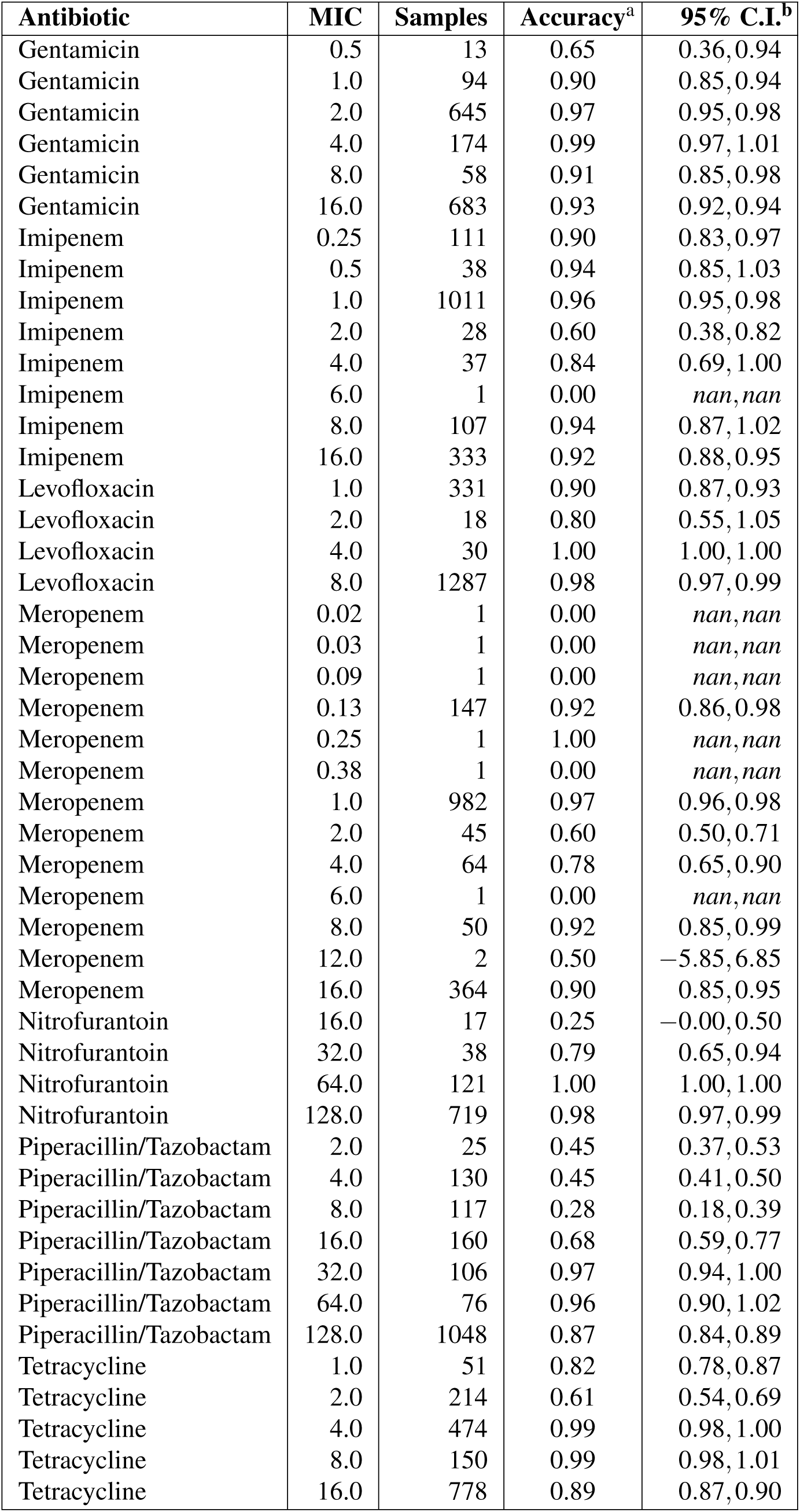
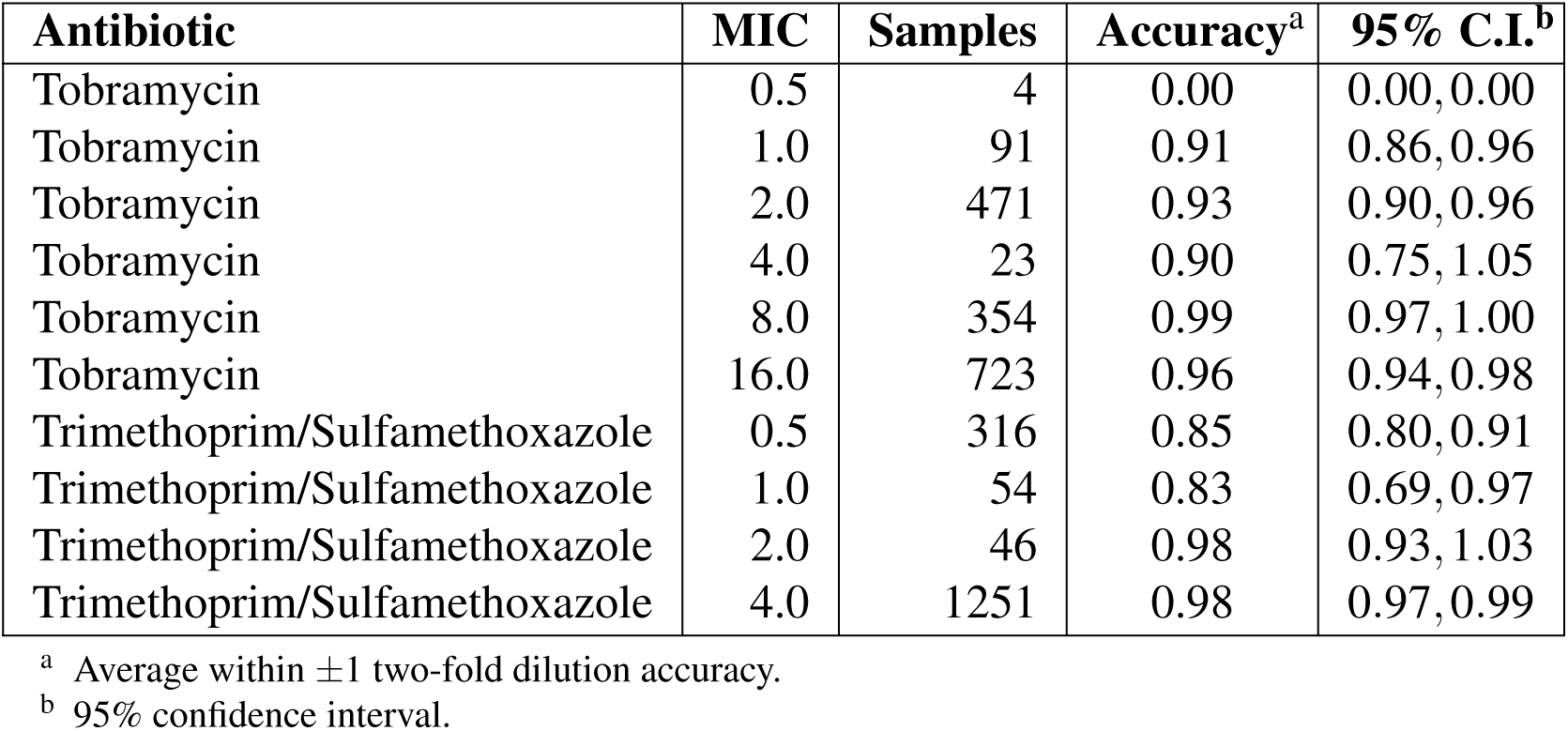
The within 1-tier accuracies for all antibiotic-MIC combinations.

**Table S4.** The within ±1-tier accuracy of the XGBoost model built from whole genome contigs, AMR genes only, and non-AMR genes respectively.

| Antibiotic | Samples | Contigs Accuracy | 95% C.I. | AMR Genes Accuracy | 95% C.I. | Non-AMR Genes Accuracy | 95% C.I. |
| --- | --- | --- | --- | --- | --- | --- | --- |
| All | 32705 | 0.92 | 0.92,0.92 | 0.92 | 0.91,0.92 | 0.92 | 0.92,0.92 |
| Amikacin | 1667 | 0.97 | 0.96,0.98 | 0.96 | 0.95,0.97 | 0.97 | 0.97,0.98 |
| Ampicillin | 1666 | 1.00 | 0.99,1.00 | 1.00 | 0.99,1.00 | 1.00 | 0.99,1.00 |
| Ampicillin/Sulbactam | 1664 | 0.99 | 0.99,1.00 | 0.99 | 0.99,1.00 | 0.99 | 0.99,1.00 |
| Aztreonam | 1644 | 0.89 | 0.89,0.90 | 0.89 | 0.88,0.90 | 0.89 | 0.89,0.90 |
| Cefazolin | 1667 | 0.96 | 0.95,0.96 | 0.95 | 0.95,0.96 | 0.96 | 0.96,0.97 |
| Cefepime | 1571 | 0.61 | 0.58,0.64 | 0.61 | 0.58,0.63 | 0.60 | 0.58,0.63 |
| Cefoxitin | 1645 | 0.90 | 0.89,0.91 | 0.89 | 0.87,0.90 | 0.90 | 0.89,0.92 |
| Ceftazidime | 1667 | 0.92 | 0.91,0.93 | 0.91 | 0.91,0.92 | 0.91 | 0.90,0.92 |
| Ceftriaxone | 1667 | 0.89 | 0.87,0.90 | 0.90 | 0.89,0.91 | 0.89 | 0.88,0.91 |
| Cefuroxime sodium | 1575 | 0.99 | 0.99,1.00 | 0.99 | 0.99,1.00 | 0.99 | 0.99,1.00 |
| Ciprofloxacin | 1664 | 0.98 | 0.97,0.98 | 0.98 | 0.97,0.98 | 0.97 | 0.97,0.98 |
| Gentamicin | 1667 | 0.95 | 0.93,0.96 | 0.96 | 0.95,0.96 | 0.95 | 0.94,0.96 |
| Imipenem | 1666 | 0.94 | 0.93,0.95 | 0.92 | 0.91,0.93 | 0.94 | 0.93,0.95 |
| Levofloxacin | 1666 | 0.97 | 0.96,0.97 | 0.97 | 0.96,0.98 | 0.96 | 0.95,0.97 |
| Meropenem | 1660 | 0.93 | 0.91,0.95 | 0.91 | 0.89,0.92 | 0.93 | 0.92,0.94 |
| Nitrofurantoin | 895 | 0.96 | 0.95,0.97 | 0.96 | 0.95,0.96 | 0.96 | 0.96,0.97 |
| Piperacillin/Tazobactam | 1662 | 0.78 | 0.77,0.79 | 0.77 | 0.75,0.80 | 0.77 | 0.76,0.79 |
| Tetracycline | 1667 | 0.89 | 0.87,0.90 | 0.90 | 0.89,0.92 | 0.89 | 0.88,0.90 |
| Tobramycin | 1666 | 0.95 | 0.94,0.96 | 0.95 | 0.94,0.96 | 0.93 | 0.92,0.95 |
| Trimethoprim/Sulfamethoxazole | 1667 | 0.95 | 0.94,0.96 | 0.94 | 0.93,0.95 | 0.95 | 0.93,0.96 |

**Table S5.** The number of different MIC values observed within the top five *K. pneumoniae* MLST types in the dataset.

| Antibiotic | MLST type |  |  |  |  |
| --- | --- | --- | --- | --- | --- |
|  | 307<br>(560 genomes) | 258<br>(404 genomes) | 16<br>(86 genomes) | 15<br>(56 genomes) | 280<br>(27 genomes) |
| Amikacin | 5 | 6 | 3 | 4 | 2 |
| Ampicillin | 1 | 1 | 1 | 1 | 1 |
| Ampicillin/Sulbactam | 2 | 3 | 3 | 2 | 1 |
| Aztreonam | 5 | 5 | 5 | 4 | 2 |
| Cefazolin | 2 | 3 | 2 | 4 | 1 |
| Cefepime | 6 | 8 | 6 | 6 | 3 |
| Cefoxitin | 4 | 4 | 4 | 4 | 4 |
| Ceftazidime | 5 | 3 | 4 | 5 | 3 |
| Ceftriaxone | 3 | 6 | 3 | 3 | 1 |
| Cefuroxime sodium | 1 | 2 | 2 | 2 | 1 |
| Ciprofloxacin | 1 | 1 | 1 | 1 | 2 |
| Gentamicin | 6 | 6 | 5 | 4 | 1 |
| Imipenem | 8 | 6 | 3 | 6 | 1 |
| Levofloxacin | 3 | 2 | 2 | 2 | 3 |
| Meropenem | 7 | 10 | 4 | 3 | 1 |
| Nitrofurantoin | 4 | 1 | 3 | 4 | 3 |
| Piperacillin/Tazobactam | 7 | 7 | 7 | 7 | 5 |
| Tetracycline | 5 | 6 | 4 | 5 | 1 |
| Tobramycin | 7 | 5 | 4 | 4 | 2 |
| Trimethoprim/Sulfamethoxazole | 5 | 5 | 3 | 4 | 1 |

**Figure S1.**
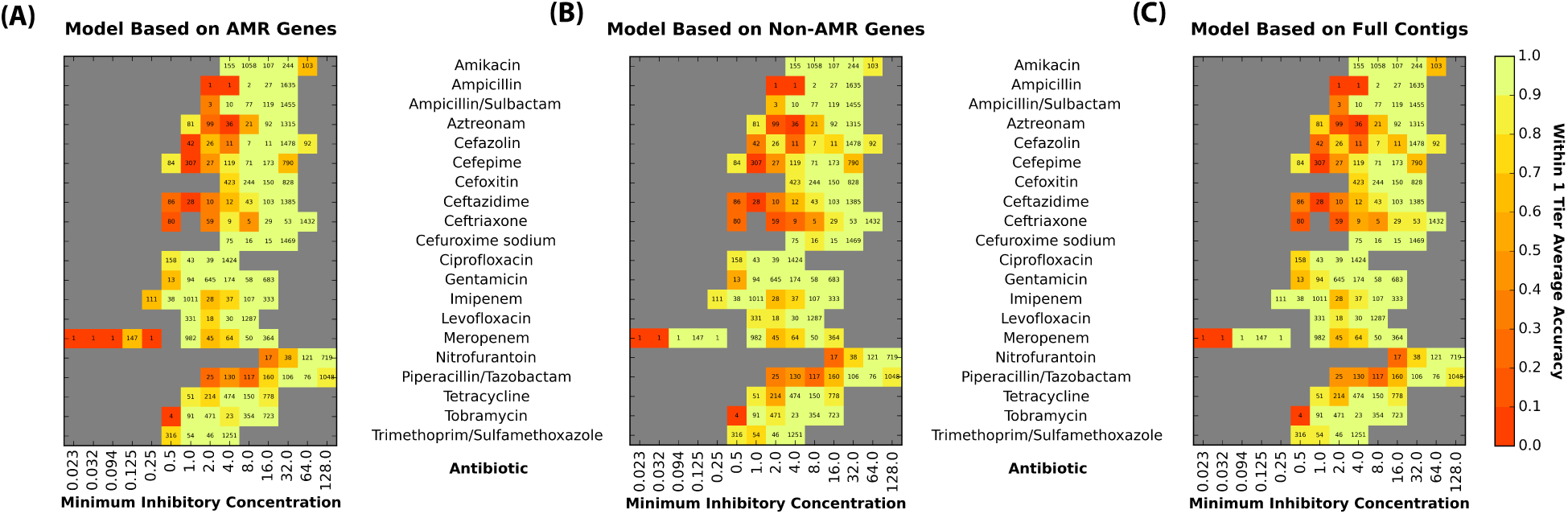
Heat maps comparing the accuracies of the XGBoost model for individual MICs generated from (A) AMR genes, (B) Non-AMR genes, and (C) full contigs. The X-axis of the heatmap shows shows the actual MIC (*μ*g/ml) for a bin and the Y-axis lists the antibiotics. The within ± 1-tier accuracy of a particular antibiotic-MIC bin is denoted by color, with red abd orange being least accurate and bright yellow and green being most accurate. The number within each cell represents the number of samples (genomes with the MIC) within the bin. The data depict genomes for which there is at least one AMR gene called by PATRIC or CARD.

